# ClustENMD: Efficient sampling of biomolecular conformational space at atomic resolution

**DOI:** 10.1101/2021.04.16.440182

**Authors:** Burak T. Kaynak, She Zhang, Ivet Bahar, Pemra Doruker

**Affiliations:** Department of Computational and Systems Biology, School of Medicine, University of Pittsburgh, Pittsburgh, PA, USA; OpenEye Scientific Software Inc., Santa Fe, NM, USA

## Abstract

**Summary:** Efficient sampling of conformational space is essential for elucidating functional/allosteric mechanisms of proteins and generating ensembles of conformers for docking applications. However, unbiased sampling is still a challenge especially for highly flexible and/or large systems. To address this challenge, we describe the new implementation of our computationally efficient algorithm ClustENMD that is integrated with ProDy and OpenMM softwares. This hybrid method performs iterative cycles of conformer generation using elastic network model (ENM) for deformations along global modes, followed by clustering and short molecular dynamics (MD) simulations. *ProDy* framework enables full automation and analysis of generated conformers and visualization of their distributions in the essential subspace.

**Availability and implementation:** ClustENMD is open-source and freely available under MIT License from https://github.com/prody/ProDy.

**Supplementary information:** Supplementary materials comprise method details, figures, table and tutorial.

## 1. Introduction

Mapping the conformational space of proteins has been a challenge, especially for large assemblies of complexes. Elastic network models (ENMs) and normal mode analysis (NMA) have proven to predict the global modes of motion of biomolecular systems, and particularly supramolecular machines in the last two decades, as shown in numerous comparisons with experimentally observed conformational changes.^1-3^ Thus, hybrid techniques combining ENM/NMA with MD have come forth as computationally efficient means for elucidating transition pathways^4-5^ and for conformational sampling for large complexes^6-7^, as we recently reviewed^8^.

ClustENM hybrid algorithm^7^ has been introduced for *unbiased sampling* of the essential subspace spanned by the softest ENM modes through integration with clustering and energy minimization of conformers. Comparison with experimental data has shown the efficiency and utility of ClustENM for investigating highly flexible proteins like calmodulin^7, 9^ as well as large assemblies such as the ribosome^7, 10-11^. More recently, ClustENM conformers have proven to facilitate protein-DNA and protein-protein ensemble docking^12^ and prediction of cryptic allosteric pockets^13^.

Such promising results have motivated us to further develop and implement the ClustENMD version in the widely used *ProDy*^14-15^ application programming interface (API) via integration with OpenMM^16^ software. This version allows us to generate more realistic conformers by performing short MD simulations even for large allosteric complexes, together with high efficiency and full automation within a Python environment.

## 2. Method and Features

The ClustENMD algorithm is explained schematically in **Figure 1A**. In *Step 1*, the input structure is subjected to ANM analysis to produce atomistic conformers using random deformations along linear combinations of a set of global ENM modes. In *Step 2*, the conformers are clustered based on their structural similarities, and a representative member is selected for each cluster. In *Step 3*, the representatives from the previous step are structurally refined by short MD simulations using OpenMM. The new conformers are then fed back to *Step 1*, each being used as a starting point for a new generation of conformers. This iterative procedure (*Steps 1-3*) is repeated for several generations to allow for sufficiently large excursions from the original energy minimum.

**Figure 1.**
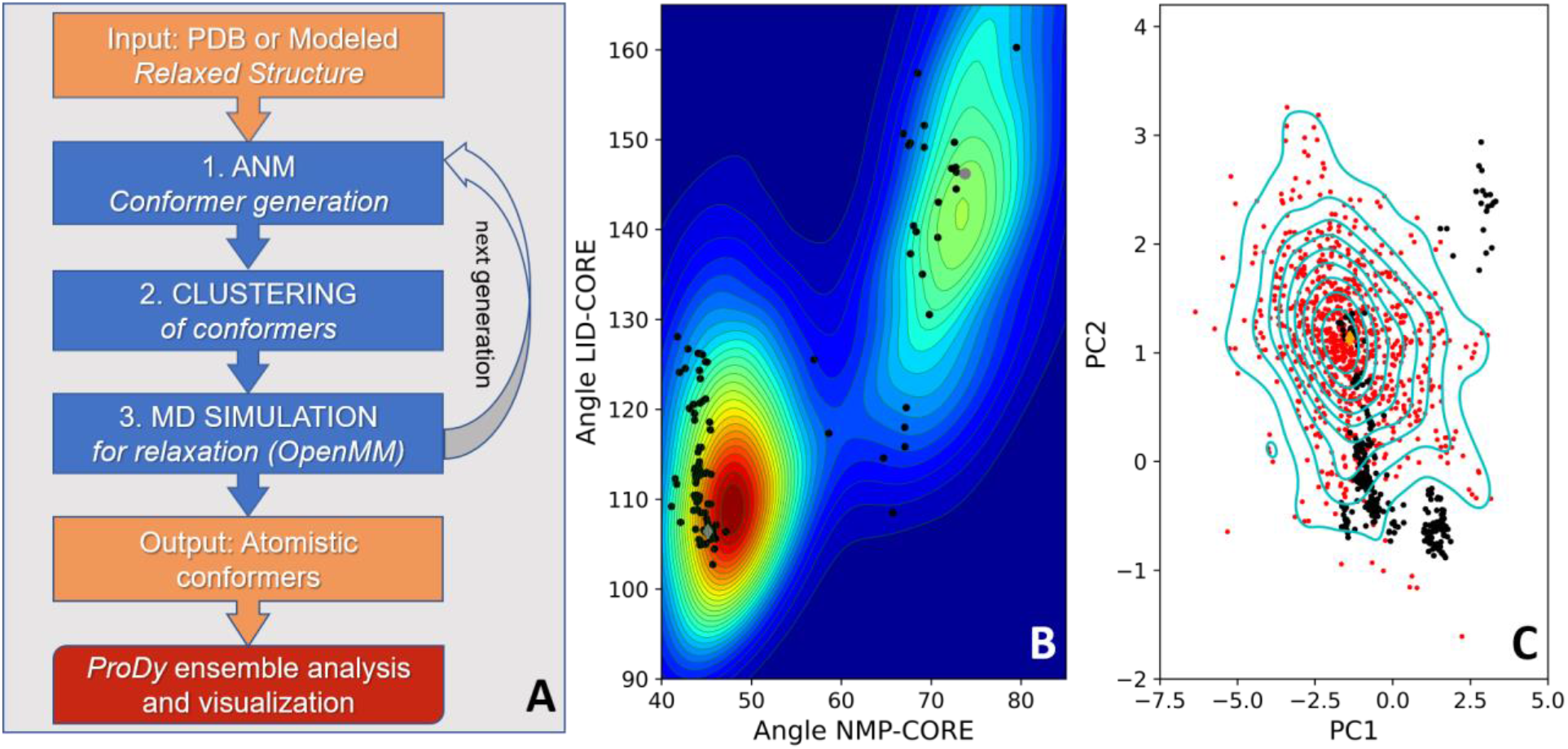
ClustENMD workflow and conformational sampling. **(A)** ClustENMD algorithm consists of three steps in each generation: (1) ANM conformer generation by ProDy, (2) hierarchical clustering of conformers, and (3) MD simulations for structural relaxation by OpenMM. **(B)** *Adenylate kinase (AK)*. Population distribution of ClustENMD conformers is plotted on the LID-Core and NMP-Core angles space of AK. This distribution contains a total of 1804 conformers from 5-generation runs, namely 3 runs starting from the open/apo (gray circle) and 3 runs from the closed/inhibitor-bound (gray diamond) x-ray structures. Homologous experimental structures are depicted as black circles. **(C)** *HIV-1 RT*. Conformational surface plotted along the first two principal components obtained from experimental structures (black circles), onto which the ClustENMD conformers (5 generations, implicit solvent) and the initial structure (orange diamond) are projected. Blue lines indicate the density levels of 903 ClustENMD conformers. The distributions of conformers in panels **B** and **C** were produced by kernel density estimate (KDE) using Gaussian kernels.

ClustENMD has the following features (see Supplementary Material and Tutorial for details):

- Implemented as a class in *ProDy*
- Integrated with OpenMM (*Step 3*)
- Applicable to multimeric complexes/assemblies, comprising protein, RNA and/or DNA chains
- Input structure either retrieved from the Protein Data Bank^17^ (PDB) or provided by the user in PDB file format
- Addition of hydrogens and any missing heavy atoms in the residues of the input structure by PDBFixer/OpenMM
- MD simulations performed by OpenMM, either in implicit solvent^18^ (Amber99SB^19^ forcefield) or explicit solvent (Amber14^20^ and TIP3P-FB^21^ forcefields) using periodic boundaries
- Anisotropic network model^22^ (ANM) for conformer generation using a set of global modes (*ProDy*)
- Pairwise root-mean-square deviation (RMSD)-based hierarchical clustering
- Atomic coordinates of conformers saved as *ProDy* ensemble, and/or in PDB/DCD format
- Analysis of output conformers using diverse *ProDy* modules, e.g. ensemble analysis
- Fully automated pipeline, from input PDB file to the generated ensemble of conformers
- High computational efficiency on GPU-architecture

## 3. Illustration

ClustENMD results for two case studies are presented in **Figure 1** using simulations in implicit solvent model and heating up (HU) the system to 300 K. **Figure 1B** presents the population distribution for adenylate kinase (AK). AK is known to undergo a large conformational change (RMSD change of 7Å) between open (apo) and closed (bound) states. The contour plot corresponding to the population distribution of ClustENMD conformers are displayed on the 2-dimensional (2D) space spanned by the two inter-domain angles, namely LID-Core and NMP-Core^23^. This plot is based on six independent runs, each comprising 5 generations (see **Table S1** for details on all systems/runs). Other homologous experimental structures retrieved from the PDB by *ProDy* are shown on the same plot (*black dots*). ClustENMD conformers sample the two states as well as the transition region between them (see **Figure S1** for runs with more generations).

**Figure 1C** displays the 2D space for hetero-dimeric HIV-1 reverse transcriptase (RT), a large enzyme (*N* = 1,000 residues). Here, the population distribution (contour plot) is projected onto the essential space of experimentally resolved structures. The axes denoted by the first two principal components (PC1 and PC2) are derived from the principal component analysis (PCA) of 365 experimental structures (*black circles*) resolved for RT under different conditions (oligonucleotide/inhibitor-bound or unbound). ClustENMD conformers (*red circles*) projected onto this space can sample the close neighborhoods of most experimental structures (see **Figure S2** for other runs including those in explicit solvent). Conformational sampling of HIV-1 protease is shown in **Figure S3**.

High efficiency of ClustENMD is reflected by the average run time of a 5-generation run generating 300 conformers (presented in **Figure 1**), which takes 8 minutes (AK, 214 residues) to 27 minutes (RT, 978 residues) on a NVIDIA® GeForce® RTX 2080 Ti graphics card.

## 4. Concluding remarks

The ClustENMD algorithm is implemented within *ProDy*^14-15^, an open-source Python API for protein structure, dynamics, and sequence analysis, containing multiple modules. The *ProDy* package (downloaded more than 2.4 million times and visited by 150,000+ unique users worldwide) ensures broad dissemination of ClustENMD to the research community in addition to providing accessory tools for analyses and visualization. The current version of ClustENMD is unique in performing unbiased sampling with high computational efficiency, augmented by fully automated and user-friendly features upon integration with *ProDy* and OpenMM.

## Supporting information

Supplementary Materials including tutorial

