## Supplementary Materials including tutorial for "ClustENMD: Efficient sampling of biomolecular conformational space at atomic resolution"

##### ClustENMD Method

We provide below more details on the ClustENMD method, its new features and implementation (see also Tutorial). A single generation ($i^{th}$ generation) of this iterative procedure comprises *Steps 1-3*, which can be repeated for as many generations (default 5) as necessary depending on the system size and flexibility.

*Input:* The starting structure can be either a PDB structure or a model provided by the user. User-provided structures should be in standard PDB format including chain ids. The input structure is parsed by *ProDy*[1-2].

After parsing, PDBFixer/OpenMM[3] adds hydrogens, as well as the missing heavy atoms in a residue that has been partially resolved. (Note that we do not use PDBFixer's option for adding residues/segments missing in the PDB structure.) Additionally, non-standard residues are changed to standard counterparts and disulfide bonds are taken care of by PDBFixer.

This fixed structure is further relaxed by OpenMM (see procedure in *Step 3*). The zeroth generation contains only this conformer in its set, which is the input to *Step 1*.

*Step 1: Conformer generation by ANM.* New conformers are generated by 'sampleModes' function in *ProDy*. For this aim, the following steps are executed on each *parent* conformer coming from the previous generation (either the input structure from the zeroth generation or all new conformers coming from *Step 3* produced in the $i^{th}$ generation):
(i) ANM is performed using a user-specified cutoff radius (default 15 Å). The coarse-grained nodes of the network correspond to CA for protein chains, C2, C4` and P for RNA/DNA chains.
(ii) Slowest *m* modes (default 3) are specified by the user to generate *n* new conformers (default 50). Note that the coarse-grained ANM modes are extended to the full atomic representation before sampling.
(iii) Deformation vectors (total number *n*) are formed by linear combinations of the modes with random coefficients, weighted by their eigenvalues.
(iv) New atomistic conformers are generated by deforming the parent conformer's coordinates along each deformation vector, ensuring that the average RMSD of new conformers with respect to the parent is equal to a user specified RMSD value (default 1 Å, may be specified differently for each generation).

The original version of ClustENM[4] performed complete enumeration of all mode combinations instead of random combinations for deforming the parent conformer. The same scheme is still available and can be also applied by choosing the version 1 (v1).

*Step 2: Clustering of new conformers.* Hierarchical clustering is performed using SciPy[5], based on pairwise RMSDs of all conformers coming from the $i^{th}$ generation (*Step 1*). Clusters are formed by specifying either the maximum number of clusters or a threshold value for RMSD, which can be set to different values for each generation. The centroid of each cluster is chosen as its representative that is carried on to the next generation.
*Step 3: Relaxation of conformers.* MD simulations are performed by OpenMM on each cluster representative (obtained in *Step 2*), either in implicit or explicit solvent with periodic boundary conditions. Relaxation comprises one or more of the following steps in sequential order:

(i) *Energy minimization (EM)*: Fast conformational search for large assemblies may be carried out by using only EM, which was the case in original implementation of ClustENM[4].
(ii) *Heating up (HU)*: Minimization may be followed by HU to the desired temperature (default 303.15 K), which is achieved incrementally.
(iii) *MD simulation*: The user can also run a short MD simulation after heating-up phase.

*Details of simulations.* The default parameters of OpenMM are as follows:

- EM is performed using 10 kJ/mol as the default tolerance for convergence.
- Time step size for MD runs is set to 2 fs by constraining all bond lengths including H atoms.
- Non-bonded interaction cutoff is fixed at 10 Å.
- Langevin integrator friction coefficient is fixed at 1 $ps^{-1}$.
- Center of mass drift is removed at each time step.
- Other system parameters that can be specified by the user with their respective default parameters given in parenthesis: temperature (default 303.15 K), pH (default neutral pH), ionic strength (default neutral) and water padding (default 1 nm) using periodic boundary.
- Explicit solvent with periodic boundary conditions is applicable for protein and RNA/DNA chains. Implicit solvent is applicable to protein chains only.

*Output:* Relaxed conformers coming from all generations, including the initial/zeroth, form the final ensemble. They can be analyzed by the 'Ensemble Analysis' and related visualization tools in *ProDy*, e.g. comparison with known structures resolved for the same protein under different conditions, or sequence homologs above a user-defined sequence identity.

**Table S1.** Simulation details of the case studies presented.

| Protein | PDB id *State* | ClustENMD[a] | | | MD runs[b] | No of conformers/structures | | Related figures |
|---|---|---|---|---|---|---|---|---|
| | | Runs | *m* | *g* | | ClustENMD[c] | Experimental[d] | |
| Adenylate kinase (AK) 214 residues | 4ake[6] *Apo* | 3 | 3 | 5 | Implicit (HU) | 903 | 109 (40% sequence identity) | 1B, S1 |
| | | 3 | | 10 | | 3302 | | S1 |
| | | 3 | | 15 | | 7199 | | S1 |
| | 1ake[7] *Complex* | 3 | 3 | 5 | | 901 | | 1B, S1 |
| | | 3 | | 10 | | 3300 | | S1 |
| | | 3 | | 15 | | 7187 | | S1 |
| HIV-1 protease 198 residues | 1tw7[8] *Apo* | 3 | 5 | 10 | Implicit (HU) | 3300 | 768 (90% sequence identity) | S3 SMov1 |
| | 1bve[9] *Complex* | 3 | 5 | 10 | | 3302 | | S3 SMov1 |
| HIV-1 RT 978 residues | 2b6a *Complex* | 3 | 3 | 5 | Implicit (HU) | 903 | 365 (90% sequence identity) | 1C |
| | | 3 | 3 | 10 | | 3301 | | S2 |
| | | 1 | 5 | 10 | Explicit (HU+15ns) | 1101 | | S2 |

[a] ClustENMD details: number of independent runs, number of slowest modes (m) and generations (g). Average RMSD between successive generations of conformers is 1 Å. The small ligand(s) that are present in the complex structures are not included during sampling/simulations.
[b] MD details: Either implicit or explicit solvent is used. HU refers to the heating up of each conformer to 300 K after energy minimization (takes about 3 ps). The last row for HIV-1 RT includes 15 ns simulation after HU.
[c] Total number of conformers generated in all independent runs.
[d] Number of experimentally resolved structures retrieved by ProDy for the investigated protein, used for comparative analyses of the predicted conformers. The corresponding minimum sequence identity percentages to retrieve the homologous structures are written in parentheses.

#### Supplementary Figures

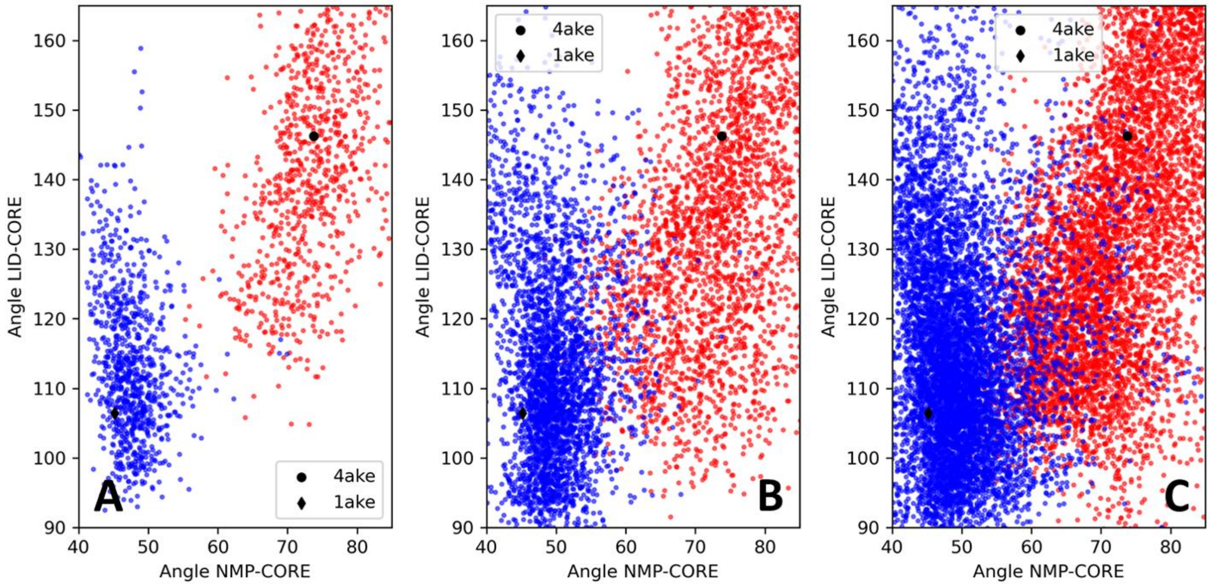


**Figure S1.** ClustENMD conformers for adenylate kinase. Conformers generated starting from the open/apo (PDB id: 4ake) and closed/complex (PDB id: 1ake) states are shown in *red* and *blue* dots, respectively. Three independent ClustENMD runs of **(A)** 5 generations (counterpart of **Fig. 1B**), **(B)** 10 generations, **(C)** 15 generations (see Table S1 for details). The transitions from closed-to-open and open-to-closed become more apparent with increasing number of generations. Conformers starting from the open (closed) structure can approach the closed (open) structure with minimum 2.9 (4.0) Å RMSD in 5 generation runs (panel **A**). Increasing the number of generations leads to even lower RMSDs (2.2 Å for open-to-closed and 2.4 Å closed-to-open in 15 generation runs, panel **C**).

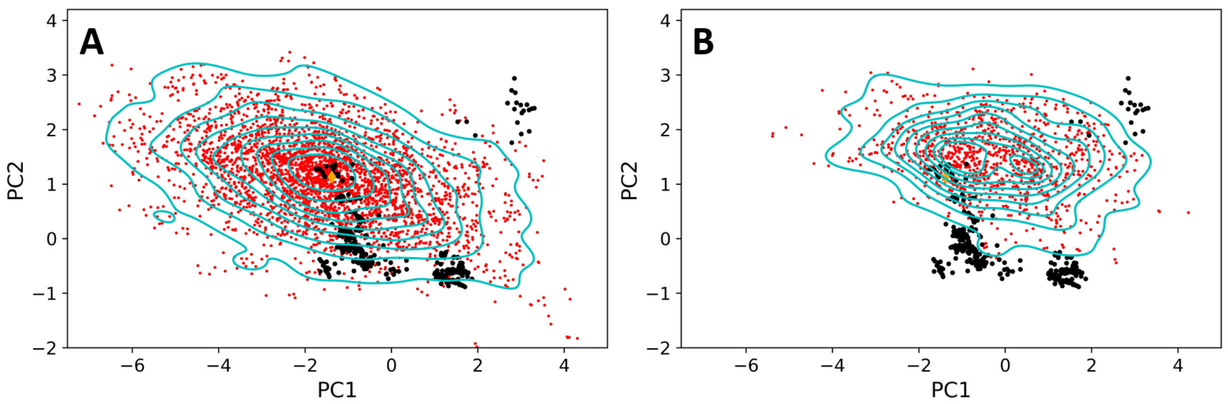


**Figure S2.** ClustENMD sampling for HIV-1 RT (initial complex, PDB id: 2b6a). **(A)** Three independent runs in implicit solvent, each carried out for 10 generations with heating-up (HU) phase only. This is the counterpart of **Fig. 1C** including 5 generations. **(B)** 10-generation run in explicit solvent with 15 ps MD following HU. Comparison of KDE plots indicates that implicit solvent simulations with HU should be preferred for computational efficiency (default in the current application) for highly flexible proteins like HIV-1 RT.

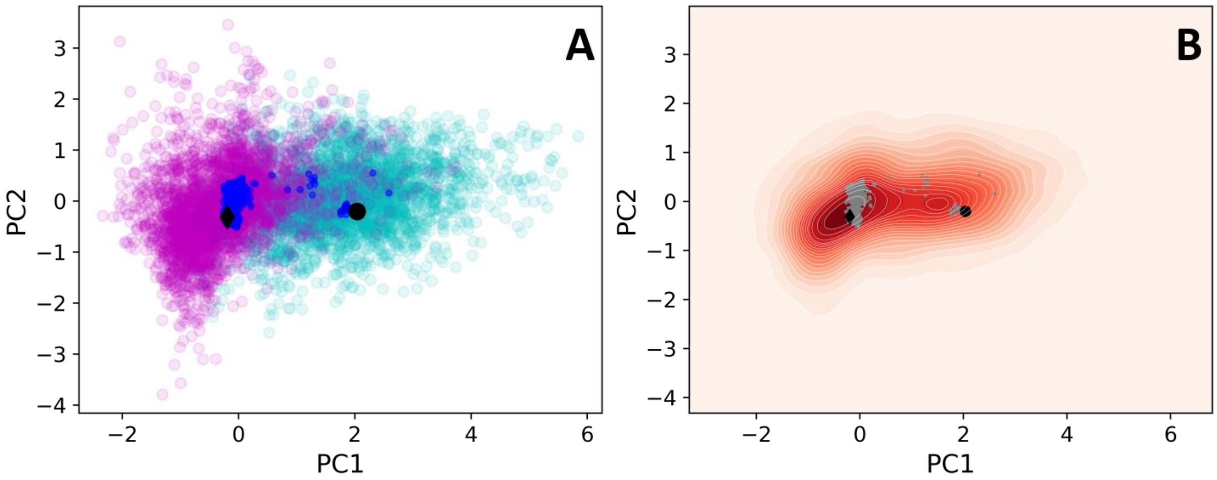


**Figure S3.** ClustENMD sampling for HIV-1 protease. **(A)** Conformers starting from the open/apo (PDB id: 1tw7) and closed/complex (PDB id:1bve) states of the protease are shown by *cyan* and *magenta circles*, respectively. The principal components are based on 768 experimental structures (*blue dots*) with the respective initial structures indicated by *black circle* and *diamond*. This distribution contains a total of 6,602 conformers from 10-generation runs (3 runs starting from the open and 3 runs from the closed states). **(B)** Density of states using the same conformers in panel **A** is represented by a KDE plot together with the experimental structures (*gray dots)*. Highly populated states correspond to darker red regions.

### ClustENMD Tutorial

by Burak T. Kaynak and Pemra Doruker

April 3, 2021

#### Introduction

ClustENM [1] is a highly efficient, unbiased conformational search algorithm. The new version of this hybrid method, ClustENMD [2], implemented in ProDy integrates random sampling (conformer generation) along global ANM modes, hierarchical clustering of generated conformers and further relaxation by MD simulations using OpenMM at each generation/cycle. Starting from a single conformer, new conformers can be generated for highly flexible and/or large biomolecular assemblies composed of proteins, RNA and/or DNA chains.

##### Required Programs

The latest versions of **ProDy**, **OpenMM**, and **PDBFixer** are required for ClustENMD.

#### Simulation and Analysis

First, we will make the necessary imports for this tutorial.

[1]:
```
import numpy as np
import matplotlib.pyplot as plt
import seaborn as sns
import pandas as pd
import prody as pr
```

##### Preparing the system and running a ClustENMD simulation

We start our calculations by parsing the structure, of which we would like to sample conformations. For this tutorial, we will fetch the X-Ray structure of HIV-1 protease in open conformation in the absence of any inhibitor (PDB id: 1tw7) from the PDB server.

It is important to note that if the starting structure is provided by the user, it should satisfy the PDB file standards, e.g., the chain IDs need to be set properly.

The pdb file (PDB id: 1tw7) is fetched by the method `parsePDB`. Please check the **ProDy Basics tutorials** for the details.

[2]:
```
ag = pr.parsePDB('1tw7', compressed=False)
```

```
@> PDB file is found in working directory (1tw7.pdb).
@> 1890 atoms and 1 coordinate set(s) were parsed in 0.03s.
```

ClustENMD is implemented as a ProDy class, named as `ClustENM`, so we can instantiate an object of it. You can provide a title, but it is optional.

[3]:
```
clustenm = pr.ClustENM()
```

Before running a simulation, we need to set the atom group that we would like to use. This method uses PDBFixer to add all hydrogen atoms as well as any missing heavy atoms in any partially resolved residues. Note that even though PDBFixer can add any residues/segments that are not resolved in the PDB structure, we are not using this option of PDBFixer. Instead, we leave modeling of those parts to the user. User-provided models should include chain IDs in their PDB files.

At this step, you can also set the pH level of the solution to select the protonation states for adding hydrogens by setting the `pH` parameter (default `pH=7.0`).

[4]:
```
clustenm.setAtoms(ag)
```

```
@> Fixing the structure ...
@> 3108 atoms and 1 coordinate set(s) were parsed in 0.03s.
@> The structure was fixed in 1.91s.
```

After setting the atoms, you can write the fixed PDB file by the method `writePDBFixed`.

[5]:
```
clustenm.writePDBFixed()
```

A ClustENMD simulation is started by the `run` method. This method accepts numerous parameters, and we will only cover the essential ones to perform a simulation in this tutorial. Therefore, we would like to encourage the readers to refer to the docstring of this method for the description of all parameters.

As this method is iterative, the user needs to set the number of generations (default `n_gens=5`). Depending on the system size, its flexibility, and the computational resources avaliable, the user can increase or decrease the number of generations. In this tutorial, we are using its default value.

The parameters regarding the main steps of the method can be grouped as follows:

1. **ANM sampling:**

   `cutoff` : Cutoff distance ($\AA$) for pairwise interactions used in ANM computations (default is 15.0).

   `n_modes` : Number of global modes for sampling (default is 3).

   `n_confs` : Number of new conformers generated from each parent conformer (default is 50).

   `rmsds` : RMSD ($\AA$) of new conformers with respect to the parent (default is 1.0).

   `v1` : Full enumeration of ANM modes, which is used in the original ClustENM method (default is False).

In the current ClustENMD version, ANM sampling is done randomly by the ProDy method `sampleModes`, where the RMSD value corresponds to the average RMSD of the new conformers with respect to the parent conformer. As the bigger RMSD value yields larger excursions from the parent, the user should be cautious on increasing its value. In contrast the original ClustENM [1] uses the full enumeration of ANM modes with fixed maximum RMSD, which can be enabled by setting `v1=True`. In both cases, we suggest using the first 3 to 5 global modes as they are known to facilitate the conformational transitions.

2. **Clustering:**

   `maxclust` : Maximum number of clusters to be formed in each generation (default is None).

   `threshold` : RMSD threshold to apply when forming clusters (default is None).

   We are using **SciPy hierarchical clustering library** to cluster the conformers in each generation. Either `maxclust` or `threshold` parameter must be specified by the user. As a guideline, we suggest to use the `maxclust` parameter. Furthermore, the parameters can be not only set to a single value across the generations, but also provided exclusive to each generation as a tuple, e.g., `maxclust=(20, 40, 60)`. Increasing the number of maximum clusters in subsequent generations allows for maximum excursion from the initial structure, thus should be prefered.

3. **Relaxation via MD simulations:**

   `temp` : Temperature at which the simulation is conducted (default is 303.15 K).

   `solvent` : Solvent model to be used. Default is `'imp'`, which corresponds to the implicit solvent model (`'amber99sbildn.xml'`, `'amber99_obc.xml'`). To choose the explicit solvent model (`'amber14-all.xml'`, `'amber14/tip3pfb.xml'`), `solvent` should be set to `'exp'`. The user may choose other force fields available in OpenMM, please see the description of `force_field` parameter. However, the default force-fields named above have only been tested in ClustENMD so far. In the current implementation of ClustENMD, implicit solvent model is applicable to protein chains only. If there are any DNA/RNA chains in your structure, ClustENMD automatically uses explicit solvent.

   `padding` : Padding distance to be used for solvation (default is 1.0 nm).

   `ionicStrength` : Total concentration of ions (both positive and negative) to add. This does not include ions that are added to neutralize the system. Default concentration is 0.0 molar.

   `tolerance` : Energy tolerance to be used for energy minimization (default is 10.0 kJ/mole).

   `maxIterations` : Maximum number of iterations to perform during energy minimization. If this is 0 (default), minimization is continued until the results converge without regard to how many iterations it takes.

   `sim` : A short MD simulation using a time step of 2.0 fs is performed if `sim=True`. Note that there is also a *heating-up phase* until the desired temperature is reached before the short MD simulation. If `sim` is set to False, only energy minizimation is performed. If only a heating-up phase is to be performed, the parameters `t_steps_i` and `t_steps_g` should be set to 0 with `sim=True` (please see below).

   `t_steps_i` : Number of simulation steps for the starting conformer, i.e. zeroth generation, (default is 1000).

`t_steps_g` : Number of simulation steps for all conformers except the starting conformer, (default is 7500). If desired, time steps for subsequent generations can be varied and given as a tuple, e.g., (3000, 5000, 7000).

`platform` : Achitecture on which the OpenMM runs (default is None). It can be chosen as `'CUDA'`, `'OpenCL'`, or `'CPU'`. For efficiency, `'CUDA'` or `'OpenCL'` is highly recommended.

We suggest to use implicit solvation and GPU platform for computational efficiency. Default parameters are highly efficient on GPU platform for proteins comprising several thousand residues. For larger assemblies, the user may prefer: (*i*) to decrease the number of clusters and/or generations, (*ii*) to perform only energy minimization with/out heating-up phase, or (*iii*) to carefully shrink the padding distance in explicit solvent.

##### Performing a simulation

In the following, we will perform a ClustENMD simulation of 5 generations using the first 3 global modes. Relaxation of conformers is carried out in implicit solvent via energy minimization followed by a heating-up phase. We are conducting the simulation on a GPU platform. Simulation details will be printed out during execution.

[6]:
```
clustenm.run(n_modes=3, n_gens=5,
             maxclust=tuple(range(20, 120, 20)),
             sim=True, solvent='imp',
             t_steps_i=0, t_steps_g=0,
             platform='CUDA')
```

```
@> Kirchhoff was built in 0.02s.
@> Generation 0 ...
@> Minimization & heating-up in generation 0 ...
@> Completed in 1.94s.
@> #-------------------/*\-------------------#
@> Generation 1 ...
@> Sampling conformers in generation 1 ...
@> Hessian was built in 0.07s.
@> 3 modes were calculated in 0.04s.
@> Parameter: rmsd = 1.00 A
@> Parameter: n_confs = 50
@> Modes are scaled by 24.611726681118544.
@> Clustering in generation 1 ...
@> Centroids were generated in 0.24s.
@> Minimization & heating-up in generation 1 ...
@> Structures were sampled in 33.37s.
@> #-------------------/*\-------------------#
@> Generation 2 ...
@> Sampling conformers in generation 2 ...
@> Hessian was built in 0.07s.
@> 3 modes were calculated in 0.08s.
@> Parameter: rmsd = 1.00 A
@> Parameter: n_confs = 50
```

```
@> Modes are scaled by 21.96801859205728.
@> Hessian was built in 0.06s.
@> 3 modes were calculated in 0.07s.
...
@> #------------------/*\------------------#
@> Generation 5 ...
@> Sampling conformers in generation 5 ...
@> Hessian was built in 0.06s.
@> 3 modes were calculated in 0.03s.
@> Parameter: rmsd = 1.00 A
@> Parameter: n_confs = 50
@> Modes are scaled by 19.25666801776903.
...
@> Clustering in generation 5 ...
@> Centroids were generated in 14.04s.
@> Minimization & heating-up in generation 5 ...
@> Structures were sampled in 174.84s.
@> #------------------/*\------------------#
@> Creating an ensemble of conformers ...
@> Ensemble was created in 0.00s.
@> All completed in 558.38s.
```

The generated conformers are stored in a ClustENM ensemble object. For future reference, the paramters set for a simualtion can be saved into a file by the method `writeParameters`:

[7]:
```
clustenm.writeParameters()
```

As ClustENM ensemble is actually a **ProDy ensemble**, we can also save it by the `saveEnsemble` method:

[8]:
```
pr.saveEnsemble(clustenm)
```

[8]:
```
'1tw7_clustenm.ens.npz'
```

We also provide a method, called `writePDB`, to write the conformers into a PDB file. The boolean parameter `single` (default is `True`) of this method controls whether the conformers are stored as models in a single PDB file, or each of them are saved as a separate PDB file.

[9]:
```
clustenm.writePDB()
```

```
@> PDB file saved as 1tw7_clustenm.pdb
```

One can also load the previously saved ensemble by

[10]:
```
saved_ensemble = pr.loadEnsemble('1tw7_clustenm.ens.npz')
```

##### Features of ClustENM ensembles

As we mentioned above, ClustENM class is derived from ProDy ensemble class, therefore the methods defined for the latter, such as `getCoordsets`, `superpose` and many more can apply to

ClustENM objects as well. All conformers in generations ($i = 1, 2, 3, \ldots$) are automatically superposed onto the initial/zeroth conformer based on $C^\alpha$-atoms during a ClustENMD simulation.

There are alternative ways of indexing the generated conformers. User can either index ClustENM object by `clustenm[3]`, which picks the 3rd conformer (presumably the 2nd conformer in the 1st generation) or equivalently with the generation number and an index as `clustenm[1, 2]`. Note that indices start from 0.

Let's check we obtain the same coordinates by two alternative methods:

[11]:
```
np.allclose(clustenm[3].getCoords(), clustenm[1, 2].getCoords())
```

[11]:
```
True
```

A ClustENM object supports slicing as well. For example, if we want to select the 3rd conformer for every generation, then we only need to specify the index of the conformer in the second slot and select all in the first slot. If the desired conformers are not available in a particular generation, then they will be skipped.

[12]:
```
clustenm[:, 3]
```

[12]:
```
<ClustENM: 1tw7_clustenm (5 conformations; 3108 atoms)>
```

We can access the corrdinates of these conformers by the `getCoordsets` method:

[13]:
```
clustenm[:, 3].getCoordsets()
```

[13]:
```
array([[[ -3.95957387,  32.35691799,  -4.37383242],
        [ -4.94566778,  32.35594469,  -4.59228821],
        [ -3.63788137,  31.46009385,  -4.70897438],
        ...,
        [ -2.37337274,  29.5071206 ,  -3.7201629 ],
        [ -1.39627789,  29.60381804,  -3.27034612],
        [ -7.98974581,  31.21050202,  -4.31887029]],

       [[ -6.89570222,  32.89490785,  -5.27764023],
        [ -7.80893237,  32.7297113 ,  -5.67617107],
        [ -6.31021832,  32.07285054,  -5.23854147],
        ...,
        [ -5.32171232,  30.53324814,  -3.46080742],
        [ -4.58778402,  30.86851485,  -2.74293152],
        [-10.41683474,  31.15561532,  -5.46381784]],

       [[ -6.3447726 ,  34.20123262,  -5.5673921 ],
        [ -7.22727328,  34.01664711,  -6.02260974],
        [ -5.82362403,  33.34645491,  -5.43376411],
        ...,
        [ -4.07602444,  31.36764316,  -4.08790043],
        [ -3.22430149,  31.72057964,  -3.52540378],
```

```
 [-10.13066977,  31.95881599,  -6.06925207]],

[[ -6.03426394,  33.17008188,  -5.2525952 ],
 [ -6.90546384,  32.76869162,  -5.56882538],
 [ -5.41631979,  32.40739972,  -5.01477094],
 ...,
 [ -4.18322255,  30.96462084,  -3.54549089],
 [ -3.39843848,  31.42003303,  -2.95973127],
 [-10.00982495,  30.65422159,  -6.45285668]],

[[ -5.90545369,  33.39176383,  -5.49324755],
 [ -6.79399411,  33.26907861,  -5.95751872],
 [ -5.56441284,  32.44150355,  -5.52143941],
 ...,
 [ -2.89975089,  29.95653924,  -5.45052765],
 [ -1.8757943 ,  30.2292032 ,  -5.24180161],
 [ -9.38759977,  30.58004821,  -5.53001208]]])
```

On the other hand, we may want to select all the conformers of a specific generation. It is then enough to set the index of the generation in the first slot and select all in the second slot.

[14]:
```
clustenm[3, :]
```

[14]:
```
<ClustENM: 1tw7_clustenm (60 conformations; 3108 atoms)>
```

##### Analysing the results

We would like to show how the computed conformers populate the conformational space as regards the essential dynamics of the structure. For this aim, we perform a principal component analysis (PCA) on the generated ensemble. Next, we will project the conformers onto the space spanned by the first two PCs, which explain the highest variance of the ensemble. This can be done using **ProDy ensemble analysis**.

We are calculating PCs based on the $C^{\alpha}$-atoms. This selection can be done directly on the ClustENM object.

[15]:
```
clustenm.select('ca')
```

[16]:
```
clustenm
```

[16]:
```
<ClustENM: 1tw7_clustenm (301 conformations; selected 198 of 3108 atoms)>
```

[17]:
```
pca_clustenm = pr.PCA()
pca_clustenm.buildCovariance(clustenm)
pca_clustenm.calcModes()
```

```
@> Covariance is calculated using 301 coordinate sets.
@> Covariance matrix calculated in 0.016746s.
```

```
@> 20 modes were calculated in 0.06s.
```

We can observe the progression of the conformers by coloring them in successive generations (from initial/zeroth to the last/fifth).

[18]:
```
colors = ['r', 'm', 'c', 'orange', 'blue', 'green']

plt.figure(dpi=300)

for i in range(1, clustenm.numGenerations() + 1):
    pr.showProjection(clustenm[i, :], pca_clustenm[:2],
                      c=colors[i], label=f'{i}')
pr.showProjection(clustenm[0, :], pca_clustenm[:2],
                  c=colors[0], label='0', marker='*', markersize=10)

plt.xlabel('PC1')
plt.ylabel('PC2')
plt.legend()

plt.tight_layout()
plt.show()
```

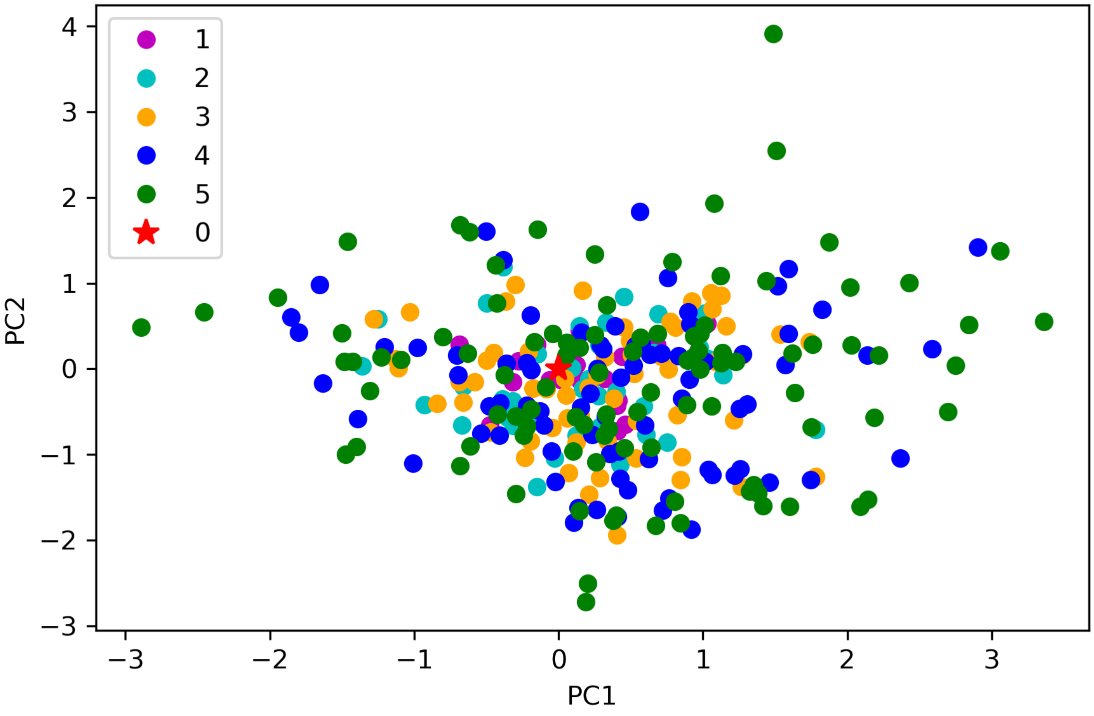


The median and maximum RMSDs with respect to the initial conformer can be calculated for the whole ensemble by

[19]:
```
rmsds = clustenm.getRMSDs()
```

[20]:
```
np.median(rmsds), np.max(rmsds)
```

[20]: (1.6681441595969058, 4.407775779940453)

One can also check the RMSDs of the conformers in each generation with respect to the initial conformer:

[21]:
```
rmsd_gens = []
for i in range(1, clustenm.numGenerations() + 1):
    tmp = pr.calcRMSD(clustenm.getCoords(), clustenm[i, :].getCoordsets())
    rmsd_gens.append([tmp.min(), tmp.mean(), tmp.max()])

df = pd.DataFrame(rmsd_gens, index=range(1, clustenm.numGenerations() + 1),
                  columns=['min', 'mean', 'max'])
```

[22]:
```
pb = df.plot.bar(color=['c', 'm', 'r'])
pb.figure.set_dpi(300)

pb.set_xlabel('Generation')
pb.set_ylabel(r'RMSD $(\AA)$')

plt.tight_layout()
plt.show()
```

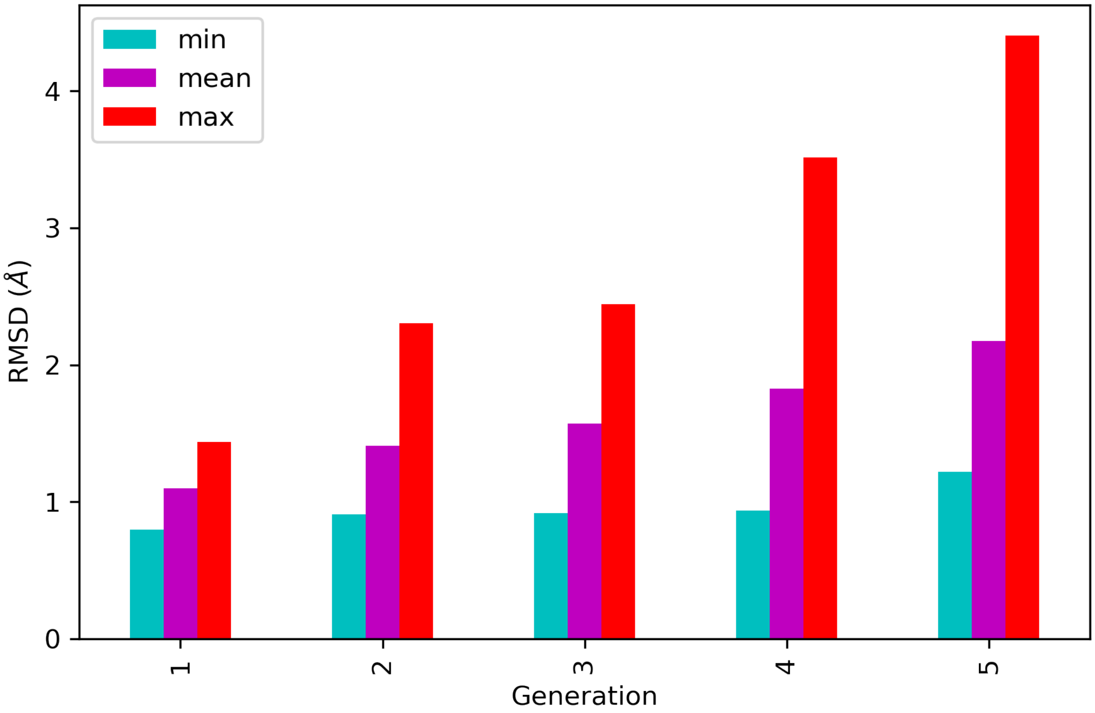

We want to also observe if our conformers approach the closed state of HIV-1 protease. For this purpose, an NMR ensemble of 28 models (PDB ID: 1bve with closed flaps) is projected onto the same subspace.

Let's first fetch these models and superpose them onto the initial/zeroth conformer. For this step, we generate an ensemble of NMR models.

[23]:
```
closed = pr.parsePDB('1bve', subset='ca', compressed=False)
```

```
@> PDB file is found in working directory (1bve.pdb).
@> 198 atoms and 28 coordinate set(s) were parsed in 0.10s.
```

[24]:
```
ens_cl = pr.Ensemble()
ens_cl.setAtoms(closed)
ens_cl.setCoords(clustenm.getCoords())
ens_cl.addCoordset(closed.getCoordsets())
ens_cl.superpose()
```

```
@> Superposition completed in 0.03 seconds.
```

At this point, we will generate the population density of the ClustENMD conformers using Seaborn implementation of kernel density estimate plot. Projected NMR conformers are also displayed as gray dots.

[25]:
```
proj_2d_clustenm = pr.calcProjection(clustenm, pca_clustenm[:2])
```

[26]:
```
plt.figure(dpi=300)

sns.kdeplot(x=proj_2d_clustenm[:, 0],
            y=proj_2d_clustenm[:, 1],
            shade=True)
pr.showProjection(clustenm[0], pca_clustenm[:2],
                  c='r', marker='*', markersize=10,
                  label='Initial')
pr.showProjection(ens_cl[2:], pca_clustenm[:2],
                  markersize=1, c='gray', label='1bve')

plt.xlabel('PC1')
plt.ylabel('PC2')
plt.legend()

plt.tight_layout()
plt.show()
```

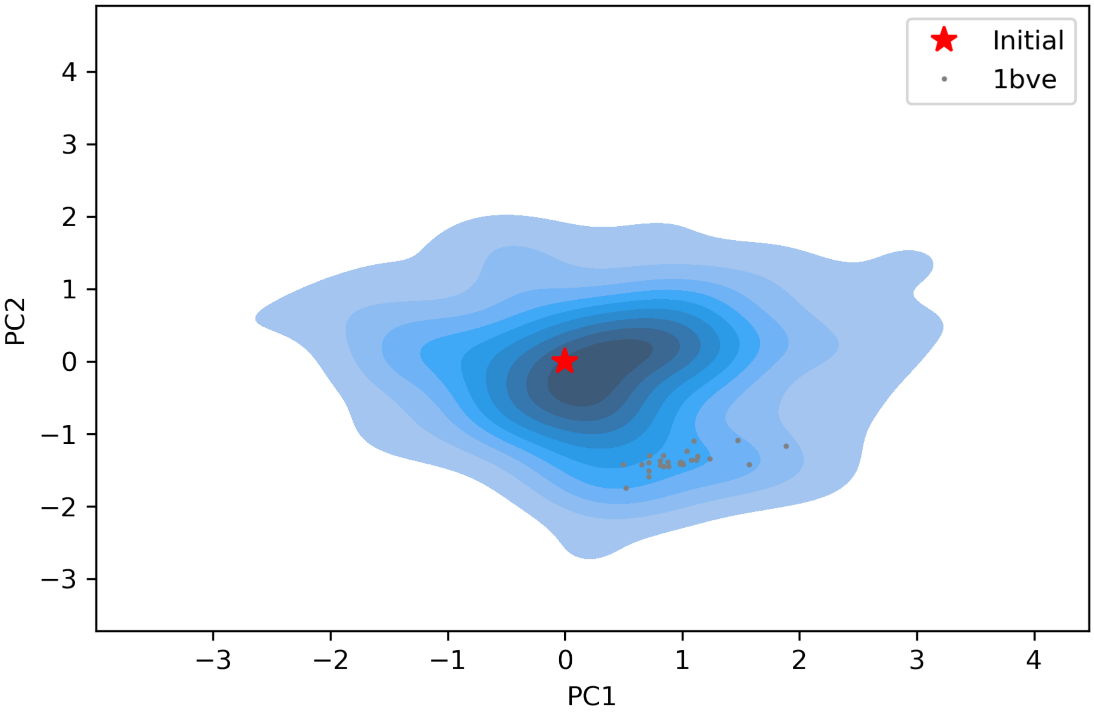


The figure above indicates that the unbiased conformer generation starting from the open state of HIV-1 protease (red star) can successfully encompass the NMR models representing its closed state (gray dots). Each time you perform a ClustENMD run, you will obtain a unique ensemble due to the random sampling and MD simulations. Therefore, it is good practice to perform at least three independent runs, and combine the resulting ensembles for analysis.

**Note:** In this tutorial we showed the variability of our generated conformers following the procedure in our original paper [1] . An alternative approach could also be followed if there are plenty of experimentally resolved homologous structures representing alternative states of a flexible protein. In the latter approach, we can perform PCA on the ensemble of experimental structures and later project the ClustENMD conformers onto the subspace defined by experimental structures (see the examples in [2]).

[2] Kaynak B.T., Zhang S., Bahar I., and Doruker P., ClustENMD: Efficient sampling of biomolecular conformational space at atomic resolution, *Bioinformatics* (submitted).
